# Inter-tool analysis of a NIST dataset for assessing baseline nucleic acid sequence screening

**DOI:** 10.1101/2025.05.30.655379

**Authors:** Tyler S. Laird, Kevin Flyangolts, Craig Bartling, Bryan T. Gemler, Jacob Beal, Tom Mitchell, Steven T. Murphy, Jens Berlips, Leonard Foner, Ryan Doughty, Felix Quintana, Michael Nute, Todd J. Treangen, Gene Godbold, Krista Ternus, Tessa Alexanian, Nicole Wheeler, Samuel P. Forry

**Affiliations:** NIST, 100 Bureau Dr. Gaithersburg, MD 20899; Aclid, 442 5th Ave #2300 New York, NY 10018; Battelle Memorial Institute, 505 King Ave, Columbus OH 43201; RTX BBN Technologies, 10 Moulton Street, Cambridge, MA 02138; SecureDNA Foundation, Aeschenplatz 6, Basel, Switzerland 4010; Department of Computer Science, Rice University, Houston, TX 77005; Department of Bioengineering, Rice University, Houston, TX 77005; Ken Kennedy Institute, Rice University, Houston, TX 77005; Signature Science, LLC; 1670 Discovery Drive, Charlottesville VA 22911 USA; Signature Science, LLC; 8501 North Mopac Expressway, Suite 100, Austin, TX 78759 USA; International Biosecurity and Biosafety Initiative for Science (IBBIS), Geneva, Switzerland; Institute of Microbiology and Infection, University of Birmingham, UK

## Abstract

Nucleic acid synthesis is a dual-use technology that can benefit fields such as biology, medicine, and information storage. However, synthetic nucleic acids could also potentially be used negligently and ultimately cause harm, or be used with malicious intent to cause harm. Thus, this technology needs to be appropriately safeguarded. Sequence screening is one component of a biosecurity protocol for preventing such harm and consists of differentiating Sequences of Concern (SOCs) from benign sequences that are not associated with pathogenicity or toxicity. There exist many fit-for-purpose tools that have been developed for DNA synthesis sequence screening. However, questions remain regarding their performance with respect to consistency of screening. To aid in determining if screening tools are harmonized in regard to baseline sequence screening, NIST constructed a test dataset based on current screening recommendations. NIST then sent blinded datasets to sequence screening tool developers for testing. Overall, there was a general agreement between the tools and NIST assignments of the sequences and all tools had a baseline performance of greater than 95% sensitivity and 97% accuracy. Disagreement on specific sequences largely arose from single tools and could be traced to differences in defining a SOC and/or methodological differences in screening algorithms.

## Introduction

Nucleic acid synthesis technology is largely considered a “dual use” technology. It can be used for the good of humanity but could have destructive potential if its use is unguarded^1^. DNA synthesis has been instrumental in understanding the basic genomic elements of life through the construction of a minimal bacterial genome^2^. It has also been used for disease diagnostics^3^, crop improvement^4^, and in biomanufacturing of biofuels^5^. Furthermore, in the future, DNA synthesis may potentially be used for storing information^6^. The capabilities of this technology, which underpins vast domains of biological research, continues to improve^7^, alongside the molecular biology tools for manipulation and assembly of synthetic DNA. As DNA synthesis becomes more accessible, however, the number of actors who could potentially misuse it to create or enhance pathogenic microorganisms grows. For example, many were unsettled by the synthesis of horsepox virus^8^, concerned that it might lower barriers to creating viruses of pandemic potential^9^.

One way to combat the potential misuse of DNA synthesis technology is for synthetic nucleic acid providers to screen DNA synthesis orders for Sequences of Concern (SOCs) that could be used to cause significant harm. In 2010, the U.S. Department of Health and Human Services (HHS) released guidelines for synthesis screening. While some of the guidelines dealt with customer screening, they also outlined details on sequence screening and record keeping. The guidelines suggested that all double-stranded DNA (dsDNA) orders (regardless of length) be screened for similarity to sequences from pathogens and toxins on the Biological Select Agents and Toxins (BSAT) list, and for international orders to also be screened for similarity to the Commerce Control List (CCL)^10^. The main goal of sequence screening is thus to distinguish Sequences of Concern (SOCs) from benign sequences. In these initial guidelines, a “best match” approach was recommended whereby synthesis providers would screen all possible 200 bp windows (and all six open reading frames) in an ordered nucleotide sequence to see if the greatest percent identity match was to a regulated agent and thus deemed a SOC. The guidelines also supported the use of screening methodologies that were equivalent or superior to the “best match” approach, along with formation of custom curated databases^10^. Identifying a sequence of concern in an order would then prompt further screening for customer legitimacy.

Around the same time (2009) as these initial U.S. guidelines were introduced, the International Gene Synthesis Consortium (IGSC) was formed with the goal of promoting both the beneficial uses of DNA synthesis technology along with appropriate biosecurity measures to mitigate potential risks^11,12^. The consortium was focused on five areas (sequence screening, customer screening, record keeping, order refusal/reporting, and regulatory compliance)^13^. In regards to sequence screening, IGSC members screen orders against a custom curated Restricted Pathogen Database as outlined in the IGSC’s Harmonized Screening Protocol^14^. Since its initial founding with five members, the IGSC has grown to include 34 members^13^.

In 2023, HHS issued updated guidelines^15^. Under the updated guidelines, the definition of an SOC is any nucleotide (≥ 200 bp) or amino acid sequence (≥ 66 aa) that is a best match to a sequence from a federally regulated agent on the Biological Select Agents and Toxin (BSAT) list or Commerce Control List (CCL), except when the same sequence is found in an unregulated organism or toxin^15^. By 2026, the guidelines suggest implementing a lower limit of 50 bp as opposed to 200 bp. Further recommendations pertaining to sequence screening methodology encourage: 1) the responsible and secure development of databases for screening SOCs that have harmful impacts on humans, animals, and plants, 2) implementing solutions for identifying sequences that pose no pathogenic or toxicity risk, 3) utilizing screening algorithms deemed equivalent or superior to the Best Match approach, and 4) developing approaches that can screen for SOCs that are split among multiple Providers, or multiple orders at a single Provider^15^.

In order to help with the development of best practices aimed at addressing the risks of nucleic acid synthesis technologies, the National Institute of Standards and Technology (NIST) began developing standards related to DNA synthesis order screening. This included the creation of datasets that could be utilized by sequence synthesis providers to demonstrate baseline sequence screening capabilities.

As DNA synthesis capabilities expand, screening requires more sophisticated computational algorithms alongside custom databases. Many Providers have thus come to rely on fit-for-purpose tools developed and maintained specifically for DNA synthesis sequence screening which can outperform more generic tools and databases^16^. Some tools currently available include Aclid^17^, The Common Mechanism^18^, FAST-NA Scanner ^19^, SeqScreen^20^, SecureDNA^21^, and UltraSEQ^22,23^. While each tool has a main goal of detecting SOCs in DNA synthesis orders, they employ different search algorithms and databases (Table 1).

**Table 1.** Description of sequence screening tools listed in alphabetical order.

| Screening Tool | Company / Organization | Algorithms Used | Databases used | Minimum screening capability | Source | Availability |
| --- | --- | --- | --- | --- | --- | --- |
| Aclid | Aclid | Sequence alignment, AI data curation | custom database | 50 nt (capable of 30 nt) | https://www.aclid.bio/ | Commercial |
| The Common Mechanism | NTI/IBBIS | BLAST, DIAMOND, HMMER, cmScan | custom biorisk/benign, NCBI nr/nt | 50 nt | https://ibbis.bio/our-work/common-mechanism/ | Open Source |
| FAST-NA Scanner | RTX BBN Technologies | Framework for Autogenerated Signature Technology for Nucleic Acids (FAST-NA), AI-resistant signatures | Custom database derived from NCBI, UniProt, PDB, and other resources | 50 nt | https://fastna.myshopify.com/ | Commercial |
| SeqScreen | Rice University / Signature Science | SeqScreen flagging logic, Bowtie2, RAPSearch2, BLASTN, ATLAS outlier detection, BLASTX (others available but not analyzed in this study = DIAMOND, Centrifuge, HMMER, MUMmer, MEGARes) | Custom BSAT, NCBI nt, Curated UniRef100, Gene Ontology (others available but not analyzed in this study = NCBI Microbial RefSeq, Pfam, REBASE, MEGARes) | 50 nt | https://gitlab.com/treangenlab/seqscreen | Open Source |
| SecureDNA | SecureDNA | “random adversarial threshold” (RAT) exact match search | Custom database including predicted functional variants | 30 nt | https://www.securedna.org/ | Open Source (proprietary database) |
| UltraSEQ | Battelle | LAMBDA2, Threat logic, K-means, AI | UniRef100, SoC protein database, curated nucleotide database | 50 nt (capable of 30 nt or less) | https://www.battelle.org/markets/health/public-health/epidemiology-and-surveillance/ultraseq | Commercial |

In the process of constructing datasets that DNA synthesis Providers could use to demonstrate baseline sequence screening, NIST sought feedback from developers of the aforementioned tools. NIST sent a blinded test dataset to four tool developers (Aclid, Battelle, IBBIS, and RTX BBN Technologies) to assess its utility and obtain feedback. Additionally, NIST ran the same dataset through two additional tools, one that was installed locally (SeqScreen^20^, created by Rice University and Signature Science) and another which utilized a locally installed client to make API calls to a remote database (SecureDNA^24^). Since the goal was to construct datasets consisting of unambiguous SOCs and benign sequences, this assessment with tool developers allowed NIST to gauge whether the industry and government interpretation of the HHS guidelines were in agreement. This manuscript discusses the construction of the test dataset and the baseline performance of the various tools using that dataset. Overall we found the dataset, as constructed, to be useful in demonstrating baseline sequence screening. Additionally, all tools performed well in identifying SOCs (i.e. they have high sensitivity) and most of the disagreed-upon sequences came about due to slightly different definitional interpretations of the HHS guidelines.

## Results

### Test Dataset Construction

A set of ∼10,000 true positive sequence fragments (each 200 bp long) were generated by starting with functionally pathogenic bacterial and viral genes from several publicly available databases. Bacterial genes were obtained from the Virulence Factor Database (VFDB)^25^, while viral genes were obtained from UniProt^26^ with the Gene Ontology term “virus-mediated perturbation of host defense response” (GO:0019049). These functionally pathogenic genes were screened to identify regulated organisms from the Biological Select Agent and Toxin (BSAT) list^27^ based on their NCBI taxonomy ID, and were aligned across all complete genomes in RefSeq^28^ to identify 200 bp gene fragments that were unique to the regulated organisms.

A set of true negative (i.e., non-SOC) bacterial and viral (phage) genomes were pulled from RefSeq and split randomly into 250, 200 bp sequence fragments. These genomes are from non-BSAT organisms that are not known pathogens. A test data set was then constructed by randomly sampling 249 bacterial true positives (TPs), 250 viral TPs, 250 bacterial true negatives (TNs), and 250 viral TNs. The 999 sequence fragments were de-identified and evaluated across six currently available sequence screening algorithms (Aclid, The Common Mechanism, FAST-NA Scanner, SeqScreen, SecureDNA, and UltraSEQ). Sequence screening results were analyzed by NIST and discussed post-hoc with the algorithm developers to understand performance.

## Test Dataset Validation

### Majority agrees with NIST dataset

We calculated aggregate metrics, taking the calls of all tools into account for a given sequence. For this, each tool was given a vote in classifying each sequence, and the majority vote for classification was compared with the NIST labels. This aggregate scoring scheme resulted in all sequences being correctly identified. Notably, there were only 4 sequences in total (0.4%) where as many as two tools disagreed with the NIST classification of a sequence (Figure 1).

**Figure 1.**
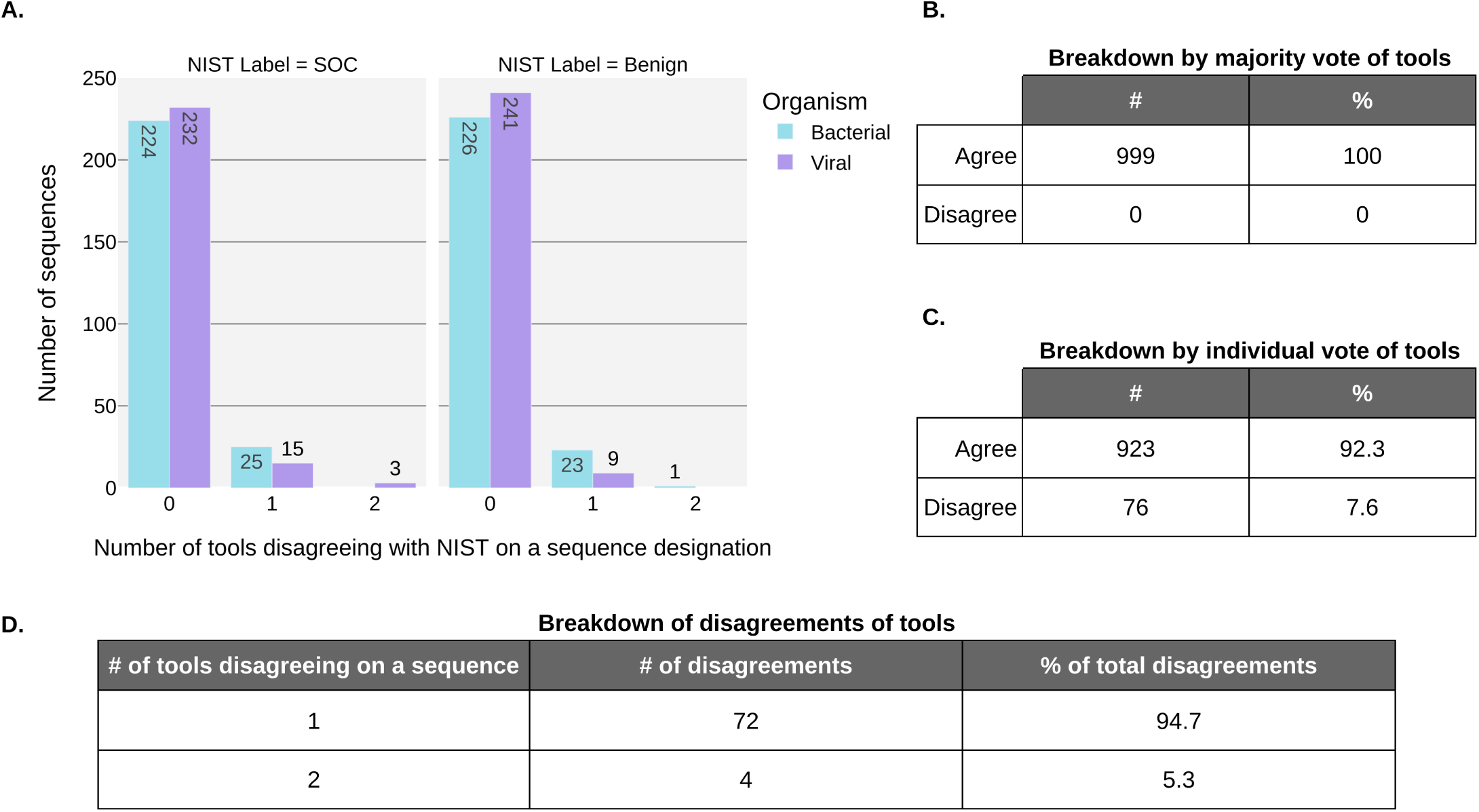
The majority of sequence classifications were in agreement with NIST. **A.** Shared x-axis represents the number of tools disagreeing with the NIST label for a sequence. A maximum of 2 of 6 total tools is shown as there were no disagreements with more than 2 tools. Plot is faceted based on sequence classification by NIST as Benign or SOC. Bars are colored based on the type of organism (bacterial or viral) from which the sequences originated. Bars are individually labeled with respective counts. **B-D.** Tables with breakdown of counts in (dis)agreement with NIST labels.

### Disagreements attributed to individual tools

In order to determine if NIST classification of SOCs and benign sequences in the test dataset aligned with calls made by tools, we compared the NIST classification of sequences with those of each individual tool. Overall, there was a general agreement between the tools and NIST assignments, as evident in Figure 1 whereby a majority of tools agreed with the label from NIST (i.e., at least 4 of 6 tools agreeing with NIST) and a majority of sequences (92.3%; see Figure 1) were unanimously identified (i.e. 0 tools disagreeing with NIST). When a tool did disagree with NIST’s designations, it usually was alone, with the other 5 tools in agreement with NIST (94.7% of disagreements), and most of these disagreements were bacterial sequences (66.7%) (Figure 1). Overall, these results point generally to subtle definitional differences between NIST and the tool developers and no widespread disagreements.

### Screening tools effectively detect SOCs

Overall, the sequence screening algorithms performed well in classifying sequences in accordance with NIST ground truth labels. We calculated the false positive rate, false negative rate, accuracy, and sensitivity for all the tools. The sensitivity and accuracy obtained by all the tools was ≥ 95% and ≥97%, respectively (Table 2). Most sequences were not disagreed upon by any tools (923/999, 92%) (Figure 1).

**Table 2.** Performance metrics of anonymized screening tools in random order.

| **Tool** | **FP Rate** | **FN Rate** | **Accuracy** | **Sensitivity** |
| --- | --- | --- | --- | --- |
| **A** | **0.004** | **0.002** | **0.997** | **0.998** |
| **B** | **0.000** | **0.034** | **0.983** | **0.966** |
| **C** | **0.022** | **0.004** | **0.987** | **0.996** |
| **D** | **0.033** | **0.002** | **0.982** | **0.998** |
| **E** | **0.006** | **0.048** | **0.973** | **0.952** |
| **F** | **0.002** | **0.002** | **0.998** | **0.998** |

### Disagreed upon sequences are attributable to varying interpretations of the HHS guidelines

Despite agreeing with a vast majority of our ground truth labels, individual screening tools differed in their disagreements with the test dataset (Figure 2). This is evident in Figure 2 which shows that disagreed-upon sequences (indicated in red) are for the most part unique to an individual tool (i.e. other rows in the column are for the most part gray).

**Figure 2.**
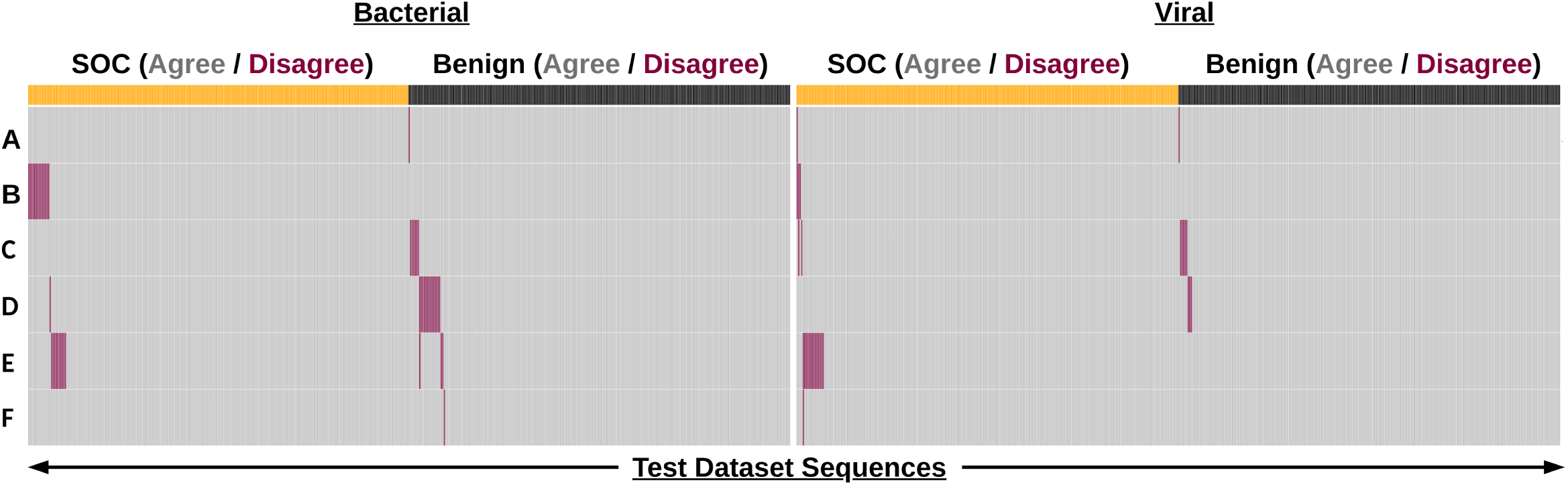
Sequence screening tools perform well on the NIST test dataset. Binary heatmap of sequence classification by anonymized tools (A-F). Each column represents a sequence (total n=999) in the dataset. Sequences are partitioned into bacterial and viral categories and labels of SOC (yellow) and Benign (black) as defined by NIST. Sequence classifications are colored depending on if they agree (gray) or disagree (red) with the NIST label.

For sequence screening, avoiding false negatives is the main priority. These represent potentially concerning sequence orders that are missed and not subjected to follow up screening. Importantly, only 43 sequences that NIST labeled as concerning were missed by one or two tools (Figure 1), and 25 of those belonged to bacteria, which is a taxonomic category where there is currently more uncertainty in the definition of risk: screening tools have been found to provide more “Undetermined” or “Optional” flagging determinations for bacterial sequences in contrast to viral sequences^29^.

While avoiding false negatives is important, preventing false positive sequences is also a key for maintaining biosecurity. While not a direct hazard, false positives can lead to additional expenses in the form of subsequent customer screening/follow-up as well as increasing screener fatigue. These expenses could in turn lead to an indirect biosecurity risk in the sense that true positive sequences may be missed due to the increased screening burden. Notably, false positives mostly occurred among bacterial sequences (Figure 1 and 2).

For the most part, the disagreements with the NIST labels (FPs/FNs) could be stratified into several categories based on differences in defining a SOC and/or methodological differences in screening algorithms. These definitional differences can be attributed to the characteristics of the sequence (nucleic acid vs. amino acid), debated sequence function (e.g., housekeeping vs. virulence), database curation (e.g., what are the sequences screened against for SOC determination and taxonomic assignment), and screening thresholds (e.g., what length of a match or how closely related does a match have to be for flagging a sequence). These definitional differences provide clarity as to why particular tools may have disagreed on a sequence, and may be resolved by ongoing standardization activities^29^.

### Nucleic acid uniqueness vs. amino acid uniqueness (false negatives)

Two sequences (#910 and #984) were disagreed upon by two tools because they had an exact match to a non-regulated organism at the amino acid level despite being unique to regulated organisms at the nucleotide level. Sequence #910 is a 200 bp fragment of the dual specificity protein phosphatase from Mpox virus (Uniprot_acc: A0A7H0DN78, nucleotide region 1:201). While it is a unique match to Mpox genomes at the nucleotide level, it has exact matches at the amino acid level to other non-regulated members of Orthopoxvirus genus such as Cowpox virus and Ectromelia virus.

Sequence #984 is a 200 bp segment of the Genome polyprotein of Swine vesicular disease virus (Uniprot_acc: A0A8F5VSD2, nucleotide region 5001:5201) which as far as our similarity searches indicate is an exact match at the nucleotide level to only Swine vesicular disease virus genomes. However, at the amino acid level, it is an exact match to many other non-regulated genomes of the Enterovirus genus. While the HHS guidelines state that an exact match to an unregulated organism is not a SOC, there is no specification on whether that is at the nucleic acid or amino acid level of a sequence.

### Definition of a virulence factor (false negatives)

While NIST curated sequences that had been annotated as a “virulence factor” (based on experimental or inferred evidence), in some areas this annotation can be ambiguous or debatable. For instance, one tool disagreed with the label of sequence #259, which was a 200 bp component of the Carbamoyl phosphate synthetase gene (*carB*) of *Francisella tularensis* (UniProt_acc: Q5NEH1, nucleotide region 1001:1201) based on identifying it as matching to a benign “housekeeping” gene. For the purposes of sequence screening, “housekeeping” genes such as ribosomal rRNAs, tRNAs, metabolic genes, etc. have generally been exempted from being flagged due to their lack of relationship to pathogenicity or toxicity. No specific definition of “housekeeping” gene has ever been provided in screening guidance, however, and so it is unsurprising to find disagreements in this area.

Likewise, the notion of a virulence factor has multiple definitions, notably including the distinction between genes that directly implement virulence functionality (e.g., toxins, invasion proteins) versus genes associated with decreased virulence when disabled, which cover a much broader range of regulatory and supporting functionality. For example, the VFDB, from which the test sequences originated, lists *carB* as a “Nutritional/Metabolic” factor involved in virulence. While mutagenesis studies do implicate *carB* in counteracting reactive oxygen species (ROS) and escaping the host phagosome^30,31^, homologs of *carB* are found widespread in bacteria, where they play an essential role in metabolism and are not associated with virulence.

While NIST gathered sequences from BSAT genomes in the VFDB, there are known shortcomings with the VFDB when trying to identify sequences of concern based on function^32^. As the definition of an SOC transitions to include functional properties, the sequences included in standardized test datasets will require further scrutiny to determine if a sequence annotated as a virulence factor is indeed a sequence of concern. Identifying essential or important genes based on mutagenesis studies does not necessarily prove that a gene is involved in pathogenesis and thus an SOC.

### Database differences (false negatives)

There were some sequences disagreed upon due to differences in databases used for screening. The definition of an SOC as defined currently by the HHS guidelines relies heavily on sequence similarity between regulated and non-regulated organisms. Identifying a sequence as an exact/best match may differ depending on what sequence database(s) a particular algorithm uses.

In constructing the test dataset NIST used only complete genomes from RefSeq to identify exact matches to virulence factor sequences. However, each screening tool includes different genomes and/or proteomes in their search databases that extend beyond what was in the NIST subset pulled from RefSeq (such as the nt or nr Genbank databases, UniRef, or even proprietary databases that are not publicly available; see Table 1). For example, sequence #372 was disagreed upon by one tool due to exact matches to non-regulated *Burkholderia* species. This sequence was a 200 bp portion of *bsaU*, a type III secretion system protein from *B. pseudomallei* (VFDB_acc: VFG002481, nucleotide region 1:201) which had exact matches to other *B. pseudomallei* and *B. mallei* genomes (both of which are regulated), as well as unclassified *Burkholderia* sp. 136(2017), sp. 129, sp. 117, and sp. 137 (which are not regulated). Despite being in RefSeq, these unclassified genomes are only assembled to the “Contig” level and thus were not included in the similarity search which would have removed them from the test dataset.

Interestingly, however, labeling the above genomes as “unclassified *Burkholderia*” is not the only valid classification. The Genome Taxonomy Database (GTDB), which classifies bacterial and archaeal genomes according to phylogenomic analysis of single-copy marker genes^33^, groups these “unclassified” genome assemblies (GCF_002900705.1, GCF_002900675.1, GCF_002900725.1, GCF_002900745.1) with *B. mallei*. Thus, differences may come about not only due to the sequences included/excluded from a tool’s database, but also from different taxonomic classifications given to the sequences. Along with the already known issues^34^, this highlights yet another challenge in using taxonomic lists for sequence screening.

### Database differences (false positives)

Most false positives were likely the result of flagging due to sequence similarity to sequences from regulated agents. Notably, the construction of the NIST database did not involve any sequence similarity searches for what were defined as true negatives. Since these sequences were obtained from genomes of non-regulated organisms, any similarity search would identify a non-regulated sequence as an exact match according to the HHS guidelines, thus categorizing it as a non-concerning sequence.

Nevertheless, these genomes from non-regulated organisms do, in some cases, have phylogenetic neighbors that are regulated. Most notably, this is the case with *E. coli* K12 in our dataset, which is closely related to regulated *E. coli* and *Shigella* species. In fact, the one benign sequence disagreed upon by 2/6 tools was sequence #582. This sequence was a genome fragment from *E. coli* K-12 belonging to the aconitase hydratase B gene (which encodes an enzyme for the conversion of citrate to isocitrate in the Krebs cycle). While the K-12 strain is a common, non-pathogenic organism used routinely in biological research, sequence #582 was presumably flagged by those tools due to its close homology to sequences from pathogenic strains of *E. coli* (STEC, EHEC, VTEC) which are CCL regulated agents^35^, although at the genetic level only the Shiga toxin gene is controlled. Among all the false positive sequences, *E. coli* K-12 sequences were the most abundantly disagreed-upon benign category, accounting for 17/33 (52% of all false positive calls).

This category also presents an example whereby consensus building activities between tool-developers resulted in a reduction in false positives (Figure 3). A prior version of one tool that predated participating in consensus building exercises (“X” in Figure 3) was used to analyze the NIST dataset. Results on this tool version showed high sensitivity to detect SOCs but lower accuracy when accounting for both SOCs and benign sequences, compared to current versions of the tools. However, after database harmonization, that tool (one of “A” through “F” in Figure 3) improved its performance on the NIST test set to obtain both sensitivity and accuracy values in line with all other evaluated tools (above 95% and 97% respectively).

**Figure 3.**
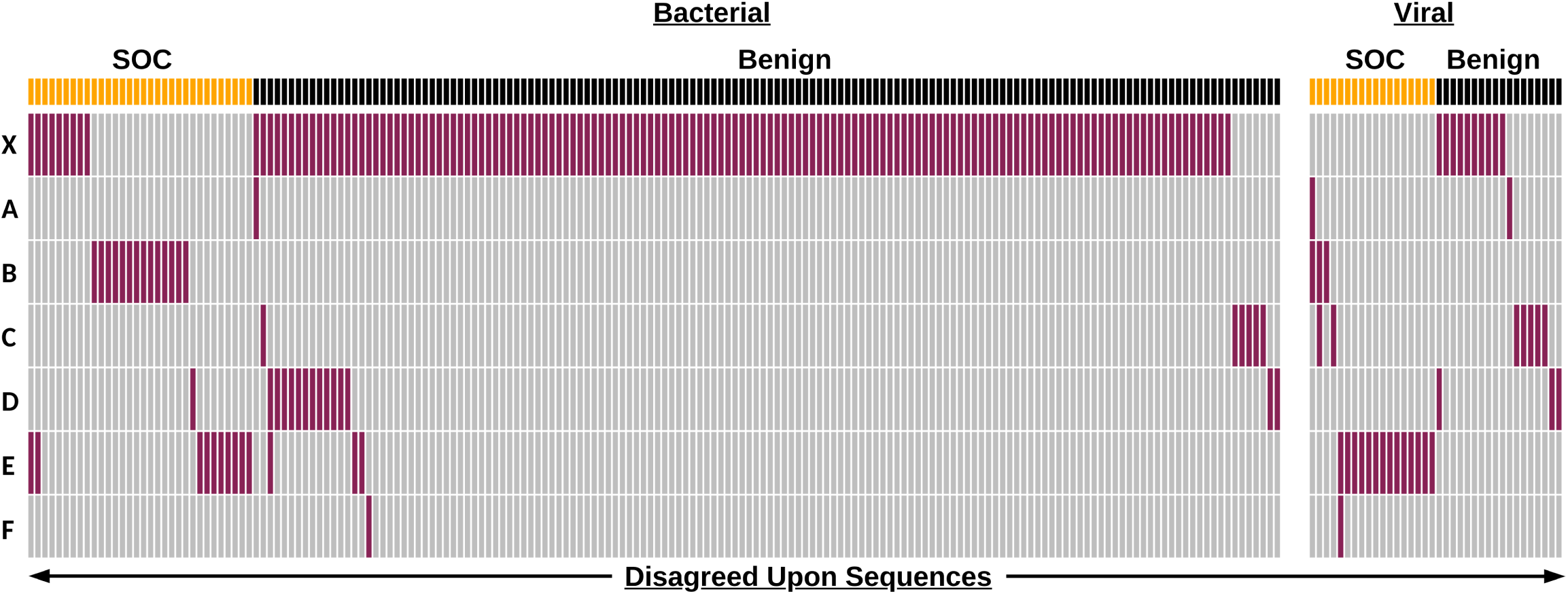
Disagreed upon sequences are primarily bacterial and reduced through consensus building. Binary heatmap of sequence classification by anonymized tools. “X” represents an earlier version of one the anonymized tools represented by “A” to “F”. Each column represents a sequence (total n=214 including “X”, n=76 for A-F only) that was misclassified. Sequences are partitioned into bacterial and viral categories and labels of SOC (yellow) and Benign (black) as defined by NIST. Sequence classifications are colored depending on if they agree (gray) or disagree (red) with the NIST label.

### More stringent screening thresholds (false positives)

While the current definition of a SOC according to the HHS guidelines relies on taxonomic similarity of sequences 200 bp or longer, the guidelines recommend a shift towards defining SOCs based on function as soon as practicable, and by 2026 screening down to a minimum length of 50 bp. Notably, certain sequences were disagreed upon by tools because they are already screening in a manner that partly aligns with this more stringent definition. For example, one tool disagreed with the NIST label for sequence #683 (a genomic fragment of Enterobacteria phage T7) because a translated 63 bp segment of it contained a hit to the structural polyprotein of Chikungunya virus (a CCL agent).

The move towards a functional definition of “sequence of concern” is important since the current definition may omit concerning sequences from both controlled and non-controlled agents. For instance, while constructing the test dataset we identified certain segments of virulence factors with exact matches to non-regulated agents. There is ongoing work related to defining a sequence of concern based on functional properties^32,36–38^. Notably, just because something is an exact match to a non-regulated organism does not mean it is benign.

### Implementation of the sequence screening assessments

Based on the results of the tool performance against the NIST test dataset, and in discussions with tool developers, key metrics were identified for a ‘passing score’ when assessing baseline sequence screening. Since it is most critical to prevent a true SOC from being synthesized without verification of customer legitimacy, the main metric of concern is sensitivity. We identified a 95% (5% false negative rate) threshold for sensitivity and a 75% threshold for accuracy (see Methods) as currently achievable by all tested algorithms. (Figure 3).

While most order streams typically contain a low frequency (<5%) of flagged sequences, we decided to have equal proportions of SOCs (TPs) and benign sequences (TNs) in this test dataset in order to better assess the ability of algorithms to identify SOCs without having to evaluate an excessively large number of TN sequences. In order to assess baseline sequence screening, adequate statistics can be achieved with 200 TP and 200 TN sequences, and additional sequences will be included that will go ungraded but serve as a way to obscure which sequences are being assessed, as well as providing a means for testing potential sequences to be graded in the future. NIST plans to generate monthly test datasets and partner with international NGOs to host and administer blinded versions of these datasets to synthesis providers.

## Conclusion

Taking into account the majority classification of sequences by current sequence screening algorithms, the NIST constructed test dataset captures the current definition of a SOC according to HHS guidelines. While there were some differences in labeling SOCs and benign sequences, these primarily localized to individual tools and appear to be due to slight definitional differences in what a tool developer or NIST identified as a SOC based on their respective interpretations of the HHS guidelines. Ongoing interactions between tool developers and shared test datasets like the one described here, provide a mechanism for harmonizing these interpretations. Notably, all of the tested tools currently screen sequences down to 50 bp and many of them implement screening against functional concerns, both of which are part of the guidelines laid out for 2026. However, future work can still be done to further harmonize differences in SOC definitions among various tools and work towards a fully comprehensive functional definition of sequences of concern.

Our analysis of baseline performance suggests that many built-for-purpose synthesis screening tools are sufficiently adept to fit into robust synthesis screening workflows. Importantly, our analysis of tool performance enabled the identification of sensitivity and accuracy thresholds for test implementation. While this study was done with tool developers, actual assessment will need to involve synthetic nucleic acid providers. While multiple providers may utilize the same tool, each tool can be run with different parameters (e.g., depending on the user or geographical location), and thus may lead to different screening outputs. Therefore, NIST test datasets will be made available for use by synthetic nucleic acid providers to assess baseline sequence screening.

## Methods

### Dataset construction

SOCs of bacterial and viral origin were obtained and labeled as “True positive” (TP) sequences in our dataset. For this, bacterial virulence factor genes from organisms on the Biological Select Agent and Toxin (BSAT) list were obtained from the Virulence Factor Database (VFDB) set A (core) nucleotide sequences. A blastn^39^ search of these nucleotide sequences against a locally constructed database of complete RefSeq genomes (n = 62,094) was performed. The results were parsed to identify 200 bp windows of nucleotides that were unique to BSAT organisms within the scope of this local database. These sequences were deemed bacterial TPs. Viral virulence factors were obtained by querying the UniProt database for entries that matched the Gene Ontology (GO) term GO:0019049 (virus-mediated perturbation of host defense response) and that belonged to one of the BSAT viral taxa (based on NCBI taxonomy id). These UniProt IDs were then mapped to genomic coding sequences in the NCBI nucleotide database (when available) and FASTA nucleotide sequences were obtained. The resulting sequences were clustered at 100% identity using the program CD-HIT^40^ to remove redundant entries that occur due to viral polyproteins. A blastn search of these nucleotide sequences was performed against the same locally constructed database of complete RefSeq genomes as above (n = 62,094). SNPs were identified on 200 bp windows with a 50 bp step size in order to obtain sequences unique to BSAT viral genomes. The pool of sequences was randomly down-sampled to 250 bacterial TPs and 250 viral TPs. While these 500 TP sequences were all tested, upon later inspection one bacterial sequence was found to be well below the 200 bp threshold and we omitted it from further analysis.

True negative (TN) sequences were obtained by randomly selecting fifty 200 bp fragments from 5 bacterial and 5 viral (phage) genomes that were from non-BSAT organisms that are not known human pathogens. Bacterial sequences came from the genomes of *Lactobacillus acidophilus* La-14, *Bifidobacterium breve* DSM 20213, *Escherichia coli* str. K-12 substr. MG1655, *Bradyrhizobium diazoefficiens* USDA 110, and *Bacillus amyloliquefaciens* IT-45. Viral sequences came from the genomes of *Escherichia* phage T7, *Inovirus* M13, *Microviridae* phi-CA82, *Leuconostoc* phage Ln-7, and *Geobacillus* phage TP-84.

The TN sequences were combined with the TP sequences in a randomized order, and given anonymized headers. The combined and anonymized dataset was then tested by each tool developer.

### Study Design

The sequences in the dataset were combined, shuffled in order, and blinded using unique numeric identifiers. This blinded dataset was then sent to the tool developers of Aclid, The Common Mechanism, FAST-NA Scanner, and UltraSEQ. Results were then sent back to NIST where they were unblinded and analyzed. Detailed analysis reports were sent to each tool developer to facilitate feedback and discussion about the utility of the dataset.

The same blinded dataset was input into one locally installed tool at NIST (SeqScreen) and another which utilized a locally installed client to make API calls to a remote database (SecureDNA). Unblinded results and analysis reports were sent to each respective tool developer similar to above.

### Tool settings

SecureDNA: The SecureDNA interface software, Synthclient, was run through the command line and individual sequence queries were sent via the available API using a Python script on 07-23-2024 with the US flagging option selected.

<u>SeqScreen</u>: SeqScreen (version 4.5) was run in a conda environment and executed to run in sensitive mode against the SeqScreen23.4 database using the command: “seqscreen --fasta SoC_test_dataset_0.2_blinded.fa --sensitive --databases SeqScreenDB_23.4 --working SeqScreen_results --threads 30”. The “flag” column was used as a binary indicator of “Flag” vs. “No Flag”.

<u>Common Mechanism</u>: The multiple columns of the Common Mechanism results file were parsed in order to identify binary “Flag” vs. “No Flag” classifications. “No Flag” sequences were defined using the query: ‘biorisk==“P” and regulated_virus == “P” and regulated_bacteria ==“P” and regulated_eukaryote==“P” and benign ==“-” and mixed_regulated_and_non_reg == “P”’.

<u>UltraSEQ</u>. UltraSEQ settings were used as described in Gemler et al^23^. Any sequence that was classified as Risk Level 1,2,3, or 4 was flagged. Risk Level 5 and 6 were considered “No Flag” for this test.

<u>Aclid and FAST-NA Scanner</u>. Aclid and FAST-NA Scanner provided results according to their default settings for current commercial deployments and reported “Flag” and “No Flag” Boolean results.

### Performance metrics

Sensitivity was defined as True Positives / (True Positives + False Negatives) where True Positives were sequences that were designated to be flagged (SOCs) in the NIST dataset. Accuracy was defined as (True Positives + True Negatives) / (Total number of sequences) where True Negatives were sequences designated as benign within the NIST dataset. The False Negative Rate was calculated as 1 – Sensitivity. The False Positive Rate was calculated as False Positives / (True Negatives + False Positives).

### Examination of disagreed upon sequences

Sequences were subjected to blastn and blastx searches using the NCBI nt/nr databases in order to identify algorithmic and database discrepancies with NIST labels. We also held follow-up conversations with each tool developer to discuss the result of each tool’s performance and better understand discrepancies.

## Author Contributions (CRediT)

- Tyler S. Laird: Conceptualization, Data curation, Formal Analysis, Investigation, Methodology, Project administration, Resources, Software, Validation, Visualization, Writing – original draft
- Kevin Flyangolts: Data curation, Investigation, Resources, Software, Writing – review & editing
- Craig Bartling: Data curation, Investigation, Resources, Software, Writing – review & editing
- Bryan T Gemler: Data curation, Investigation, Resources, Software, Writing – review & editing
- Jacob Beal: Data curation, Investigation, Resources, Software, Writing – review & editing
- Tom Mitchell: Data curation, Investigation, Resources, Software, Writing – review & editing
- Steven T Murphy: Data curation, Investigation, Resources, Software, Writing – review & editing
- Jens Berlips: Data curation, Investigation, Resources, Software, Writing – review & editing
- Leonard Foner: Data curation, Investigation, Resources, Software, Writing – review & editing
- Ryan Doughty: Data curation, Investigation, Resources, Software, Writing – review & editing
- Felix Quintana: Data curation, Investigation, Resources, Software, Writing – review & editing
- Michael Nute: Data curation, Investigation, Resources, Software, Writing – review & editing
- Todd J. Treangen: Data curation, Investigation, Resources, Software, Writing – review & editing
- Gene Godbold: Data curation, Investigation, Resources, Software, Writing – review & editing
- Krista Ternus: Data curation, Investigation, Resources, Software, Writing – review & editing
- Tessa Alexanian: Data curation, Investigation, Resources, Software, Writing – review & editing
- Nicole Wheeler: Data curation, Investigation, Resources, Software, Writing – review & editing
- Samuel P. Forry: Conceptualization, Formal Analysis, Investigation, Methodology, Project administration, Supervision, Visualization, Writing – original draft

## Competing Interests

K.F., C.B., B.G., J.Beal, T.M., S.T.M., J.Berlips, L.F, R.D., F.Q., M.N., T.J.T., G.G., K.T., T.A., and N.W. are affiliated with institutions that build and deploy biosecurity screening software.

## Funding

Work by K.F., C.B., B.G., J.Beal, T.M., S.T.M., T.A., and N.W. was partially supported by funding from Sentinel Bio. R.D. is supported in part by funds from the National Institutes of Health (P01-AI152999) and a training fellowship from the Gulf Coast Consortia, on the NLM Training Program in Biomedical Informatics & Data Science (T15LM007093). F.Q is supported in part by funds from the National Institutes of Health (GCID U19 AI144297-05). M.N is supported in part by funds from National Institutes of Health (P01-AI152999). T.J.T is supported in part by funds from the National Science Foundation (IIS-2239114), National Institutes of Health (P01-AI152999, GCID U19 AI144297-05)

## Acknowledgements

We thank Becky Mackelprang for providing valuable feedback on this manuscript

## Disclaimers

This document does not contain technology or technical data controlled under either U.S. International Traffic in Arms Regulation or U.S. Export Administration Regulations.

Certain equipment, instruments, software, or materials are identified in this paper in order to specify the experimental procedure adequately. Such identification is not intended to imply recommendation or endorsement of any product or service by NIST, nor is it intended to imply that the materials or equipment identified are necessarily the best available for the purpose.

## Data Availability

The test dataset used in this analysis is available at: https://data.nist.gov/od/id/mds2-3787

## Notes

https://data.nist.gov/od/id/mds2-3787

